# Hemispheric lateralization of Heschl’s gyrus and amygdala subdivisions are involved in differentiating happy and sad acoustic emotion

**DOI:** 10.1101/201954

**Authors:** Francis A. M. Manno, Juan Fernández-Ruíz, Fernando A. Barrios

## Abstract

Emotion is known to activate different areas of the cortex; nevertheless, how activation represents the variety of emotion is unknown. We asked whether functional activation in Heschl’s gyrus and amygdala subdivisions could differentiate happy and sad acoustic stimuli. A sparse sampling fMRI paradigm assessing hemispheric laterality by subdivision was employed to investigate this question. Differences between left Heschl’s gyrus subdivisions Te1.1 and Te1.0 compared to Te1.2 were significant for sad acoustic stimuli and right Heschl’s gyrus subdivisions Te1.0 different than Te1.1 and Te1.2 for sad acoustic stimuli, with no subdivision specificity for happy greater than sad. A prominent right greater than left difference was notable for centromedial, laterobasal, and superficial amygdala subdivisions for sad, while happy was more bilaterally activated. Activation and deactivation profiles of Heschl’s gyrus and amygdala subdivisions added to the explanation of left/right asymmetry by emotion with activation profiles significant for Heschl’s gyrus and amygdala response to sad.

## Introduction

Music elicits emotions such as sadness and happiness [1], in addition to responses that are difficult to explain such as chills [2]. Nevertheless, the psychophysiological and neural characteristics that determine what makes sound seem sad or happy have yet to be fully decoded [3, 4]. Psychophysical studies suggest that tonal properties such as frequency and amplitude give emotion to sound [5, 6]. Neuroimaging studies of happiness and sadness give credit to this hypothesis, where the difference in perception between happy or sad acoustic emotion is based on the activation of specific regions [1, 7–11]. For example, unexpected chordal sequences have elicited differential activity in the bilateral amygdala [9]. The addition of lyrics to happy or sad music alters activation in the left amygdala depending on the underlying emotion, where sad music containing lyrics demonstrates greater activation than sad music without lyrics [7]. A recent review has illustrated the importance of specific subnuclei/subdivision(s) and laterality in amygdala activation pertaining to the perception of emotion in music [3, 12], although there is controversy over the extent to which hemispheric lateralization underpins emotion perception [13–15]. Our overarching question was to explore if hemispheric laterality could differentiate emotion within Heschl’s gyrus and amygdala subdivisions.

Our aim for the present experiment was to determine if hemispheric lateralization of amygdala and Heschl’s gyrus (HG) subdivisions could differentiate a well-established acoustic emotion identification task [16, 17]. Our thinking was based on the Zatorre group principle that the right hemisphere would be more specialized for processing spectral information such as fine spectral cues while the left hemisphere would favor temporal cues [18], originally proposed by Lackner & Teuber, 1973 [19]. We hypothesized the acoustic cues in emotion would contain in general more spectral variation and would specifically lateralize right/left due to encoding happy emotion in a major key with rapid pitch changes and sad emotion in a minor key with slow undulations. We thought encoding these rapid pitch changes and slow undulations would be lateralized by the right and left hemisphere, respectively. Therefore a piece encoding happy emotion with rapid pitch changes would favor a right hemispheric lateralization activation due to the processing of fine spectral information and a piece encoding sad emotion with slow undulations would favor a left hemispheric lateralization activation due to the processing temporal cues. We further thought that along the posterior-to-anterior gradient, Te1.1 to Te1.0 to Te1.2, effectively posteromedial, to middle to anteriolateral, there would be a shift in the pitch-sensitivity of voxels to our acoustic emotion [20] possibly reflected along these axes in other subdivisions such as the amygdala. Here possibly, a ventral versus dorsal stream would be encoded by HG subdivision such that spectrotemporal features pertaining to “what” object and spatial pertaining to “where” location [21,22] would be differentially active along the Te1.1 to Te1.0 to Te1.2, gradient.

To examine this hypothesis we used an optimized sparse sampling region-of-interest (ROI) paradigm to determine whether observed differences would be elicited depending on happy and sad acoustic emotion. We postulated happy and sad emotion would be differentiated in HG and amygdala based on activation and deactivation patterns. The present study concentrated on these two ROI as opposed to a global fMRI analysis [23], because these two regions have been most commonly involved in auditory emotion [3, 12–15]. We theorized a right-leaning hemispheric activation would underpin laterality within these ROI [12–15]. We asked whether laterality and subdivision changes in activation versus deactivation could account for differences in happy and sad emotions based on an ROI analysis of lateralizing emotion.

## Methods

### Study participants

The study consisted of 12 self-reporting right-handed volunteers whom indicated no sole use of left hand, age 27.11 ± 3.20 years (range: 18 to 32 years; 6 females). All volunteers gave informed consent and were free of contraindications for psychoacoustic-MRI scanning. Participants had no musical training and were recruited from a local public university. The research protocol was approved by the Ethics Committee of the Instituto de Neurobiología at the Universidad Nacional Autónoma de México in accordance with the Declaration of Helsinki, 1964.

### Acoustic stimuli

Instrumental excerpts of classical piano known to evoke categorically a sense of sadness or happiness were taken from a previous study and curtailed to 3 seconds [MIDI files, http://www.brams.umontreal.ca/peretz; 17]. Stimuli were presented during fMRI scanning utilizing MATLAB (Statistics Toolbox Release 2012b, The MathWorks, Inc., Natick, Massachusetts) with the Psychophysics Toolbox extension [http://psychtoolbox.org/] on a HP pc (Intel Core i5-4210U CPU at 2.40GHz) with a RealTek High Definition Audio card. Stimuli were delivered via MRI-compatible headphones (AudioSystem, Nordic NeuroLab) and sound level adjustment was performed upon individuals entering the MRI unit.

### Image acquisition

Images were acquired on a 3T MR750 scanner with a 32-channel coil using parallel imaging with an acceleration factor of 2 (General Electric, Waukesha, Wisconsin). Acquisition was bottom-up interleaved. A FSPGR BRAVO structural image was acquired for co-registration with functional volumes. The structural image was a 3D T1-weighted, resolution of 1 × 1 × 1 mm^3^, FOV 25.6, slice thickness 1, TR = 8.156 s, TE = 3.18 ms, TI 450 ms, flip angle 12. The sparse sampling functional MRI acquisition consisted of 74 volumes integrated with 34 slices (3 mm thick), acquired with a single shot GE-EPI with the following parameters: TR = 15000 ms, TE = 30 ms, flip angle 90, FOV = 256 × 256 mm2, matrix = 128 × 128 (yielding a voxel size = 2 × 2 × 3 mm^3^).

### Experimental design and fMRI stimuli presentation

The order of scans acquired was first localization followed by: 1) first fMRI run, a gradient-echo, echo planar imaging sequence (GE-EPI), with left button presses for indicating sad and right button presses for indicating happy, 2) a sagittal T1 weighted Fast SPoiled GRASS (FSPGR) BRAin VOlume (BRAVO) structural image, and 3) the second fMRI run GE-EPI, with left button presses for indicating happy and right button presses for indicating sad. The first fMRI run responses were effectively counterbalanced by the second fMRI responses. The effects of scanner noise on auditory processing has been well documented; therefore, we used a sparse sampling paradigm for stimuli presentation [24]. After optimization, the protocol was 74 blocks, with 40 sound stimuli presentations, and 34 silent periods (45.95% silent blocks), for a total run time of 18 minutes and 30 seconds. The finalized functional protocol consisted of collecting one volume separated from stimuli by approximately 100 ms followed by ≈ 10.00 seconds of silence. Alike stimuli blocks (sad or happy stimuli) within an fMRI run were aggregated together and whole head statistical maps were derived.

### Functional image processing and ROI segmentation

Image processing used FSL tools (fMRIB, University of Oxford, UK) using FEAT (FMRI Expert Analysis Tool) version 5.98. The general linear model (GLM) was used to assess the relationship between the categories (i.e., EVs/regressors - happy or sad), the dependent variable (i.e., the BOLD signal), and six motion regressors (three rotation and three translation, using the double gamma function convolved with the hemodynamic response function). Higher-level analyses were performed using FLAME; in detail, single-subject first level analysis was performed using a design matrix consisting of 2 explanatory variables (EV’s) representing each of the categories (i.e. happy and sad). A fixed-effects analysis was performed to average category scans between a single-subject first fMRI run and second fMRI run. A mixed-effects higher level analysis was performed to average response categories across subjects (n=12). Correction for multiple comparisons was carried out using random field theory (voxel z > 2.3, cluster p < 0.05). Functional volumes were preprocessed for motion correction, linear trend removal, spatial smoothing using a 5 mm FWHM Gaussian kernel, and elimination of low-frequency drifts using a temporal high-pass filter with a cutoff of 100s. Preprocessing of the fMRI statistical maps included spatial realignment, coregistration with anatomical data using FSL FLIRT, and spatial normalization and alignment with MNI 152 T1-weighted MRI scans. Further image analysis was performed using custom scripts in Matlab using region of interest (ROI) segmentation from the Jülich histological atlas [25,26,] to derive 95% confidence interval probability maps associated with subnuclei specifically for the left and right hemisphere. The left and right hemisphere ROI probability maps for delineating auditory cortex were Te1.0 middle HG, Te1.1 posteromedial HG, and Te1.2 anterolateral HG; 26) and for delineating amygdala were laterobasal (LB), centromedial (CM), and superficial (SP; 25). Only areas within an ROI of activation and deactivation of at least > 100 voxels were included.

### Functional modulation analysis

Hemispheric laterality of emotion was determined using a hemispheric lateralization index to assess activation and deactivation independently for the left and right hemispheres [27]. The hemispheric lateralization index was restricted by t-value change associated with happy and sad stimuli in the absolute positive range (eg. 0 to positive numbers, effectively activation herein) and negative range (eg. negative numbers to 0, effectively deactivation herein). The nifti individual maps were coregistered to a symmetrical MNI atlas using fsl's ApplyXFM with a trilinear interpolation transformation with 6 degrees of freedom (1 rotation + 2 translation + 2 scale + 1 skew). The output map was a nii functional 4-D map in MNI symmetrical space. Where ROI probability maps pertain to asymmetrical areas of left and right hemisphere [25, 26], the symmetrical trilinear transformation accounts for those differences by the following; original right (Rorg) and original left (Lorg) for a symmetrical side (ss) is in a comparison [Rorg] – [Lorg] = Rss and [Lorg] – [Rorg] = Lss ([] denoting absolute), were we compare Rss and Lss symmetrical counterparts in the final ROI extraction of [Rss] and [Lss] by *left* side Te1.0, Te1.1, Te1.2 and *right* side Te1.0, Te1.1, Te1.2 and *left* side LB, CM, SP and *right* side LB, CM, SP [27].

The ROI statistical maps made symmetrical were restricted in the positive or negative range and also co-registered to a symmetrical MNI template, allowing direct left versus right voxel comparisons. The hemispheric activation laterality calculations reflect [R]-[L], where R and L are right and left hemispheres, respectively. Here the resultant calculation is a positive t-value reflecting right hemisphere sided laterality and a negative t-value reflecting left hemisphere laterality. The hemispheric deactivation laterality calculations reflect [-R]-[-L], with numbers in the negative range reflecting the opposite observation; here a negative number was a right hemisphere-sided laterality and a positive number was a left hemisphere-sided laterality. The hemispheric laterality calculations were separated into groups in order to distinguish the value (activation and deactivation), magnitude of lateralization, and direction (left versus right hemisphere) separately.

## Results

Laterality differences were notable for happy and sad responses within HG (Fig. 1a). Left subdivisions Te1.1 and Te1.0 were significantly different from Te1.2 in responding to sad (one sample t-test: *P* =0.0489, t = 12.99) and right subdivision Te1.0 was different from Te1.1 and Te1.2 in responding to sad (one sample t-test: *P* =0.0466, t = 13.63). Happy did not elicit greater left versus right activation for any subdivision. Differences were not notable between subdivisions within the left amygdala (Fig. 1b); however the right amygdala was significantly different in activation between sad and happy with greater activation for sad within the centromedial, laterobasal, and superficial subdivisions (one sample t-test: *P* =0.0084, t = 10.87). For sad but not happy, there was a significant difference in activation (paired t-test: *P* =0.0174, *T* = 7.4886 and *P* = 0.0924, *T* = 3.0581), between the left and right hemisphere (paired t-test: *P* =0.0211, *T* = 3.315).

**Figure 1.**
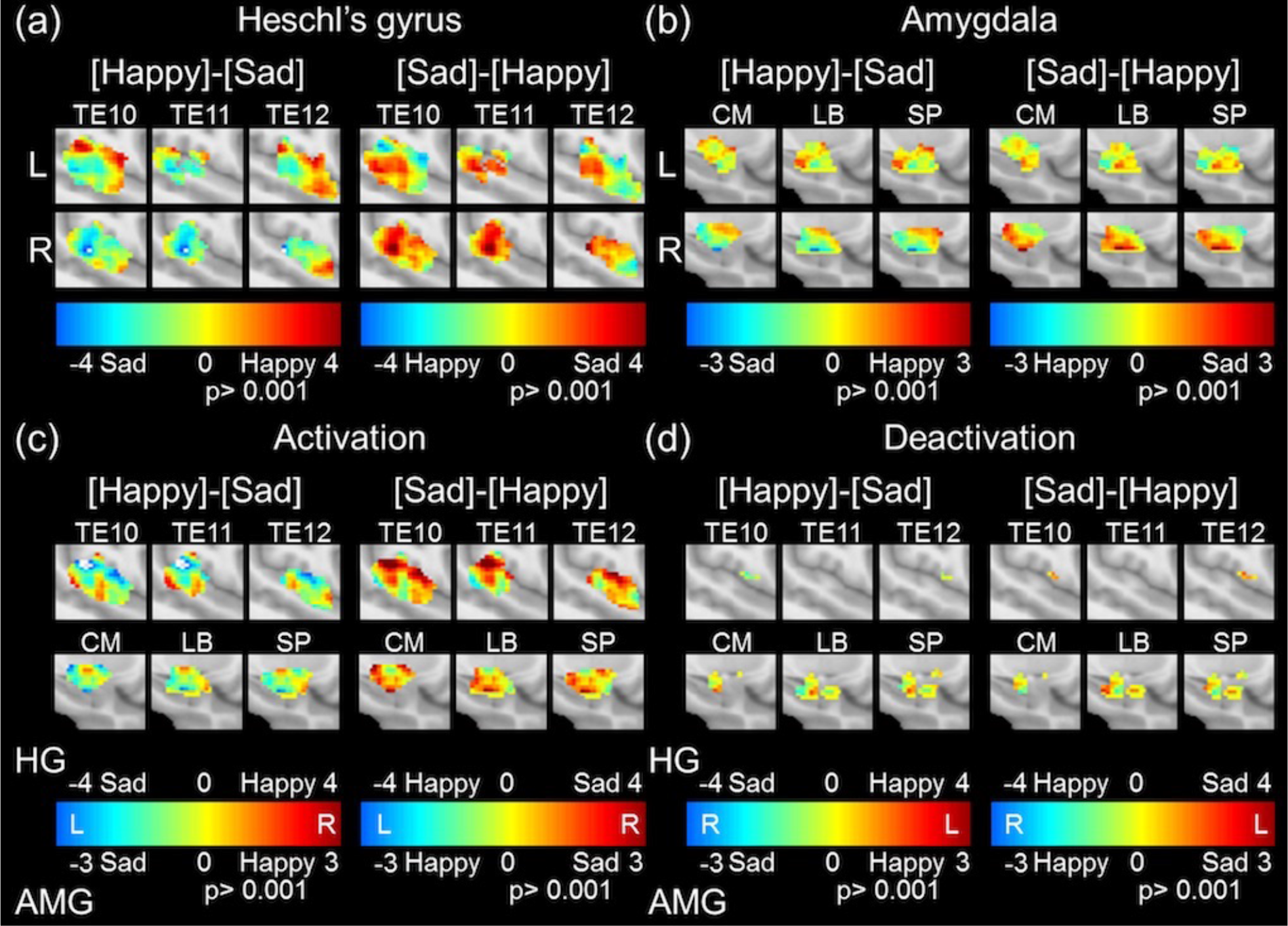
Heschl’s gyrus and amygdala hemispheric lateralization by activation and deactivation. (A) Heschl’s gyrus and (B) amygdala left (top row) and right (bottom row) for happy > sad and sad > happy by subdivision (column) for each ROI respectively. (c) Activation and (d) deactivation profiles for HG (top row) and amygdala (bottom row) for happy > sad and sad > happy by subdivision (column) for each ROI respectively. Note the color bars in (c) and (d) for HG (top) and amygdala (bottom) are different and for the activation and deactivation profiles and reveal laterality extent associated with emotion (sad and happy).

Activation and deactivation profiles revealed an interesting background to the left/right asymmetry findings (Fig 1 and 2). Activation of HG was significantly greater for sad within Te1.2 compared to the greater response for happy within Te1.1 (*P* =0.0116, *T* =3.0233; Fig. 1c upper row). As can be visualized by Fig 2a and 2b this was most likely due to a more uniformly greater response to sad over happy for the left and right hemisphere by subdivision (Fig 2c and 2d). Deactivation of HG was non-significant with Te1.0 and Te1.2 leaning toward sad while Te1.1 leaned toward happy (Fig. 1d upper row). Activation of the amygdala was significantly greater for sad within centromedial, laterobasal, and superficial subdivisions compared to happy (one sample: *P* < 0.0001, *T* = 87.04). There was no difference between amygdala subdivisions in deactivation profiles (Fig. 1c and 1d bottom row). Amygdala deactivation did not reach significance for differences between happy or sad (i.e. was robustly active responding to happy and sad emotion; Fig 2e and 2f); however, each subdivision activated more in response to sad than to happy, with right greater than left activity (Fig 2h and 2i), but responses did not reach significance due to heterogeneity. To summarize, amygdala activation for happy was more bilateral than activation for sad.

**Figure 2.**
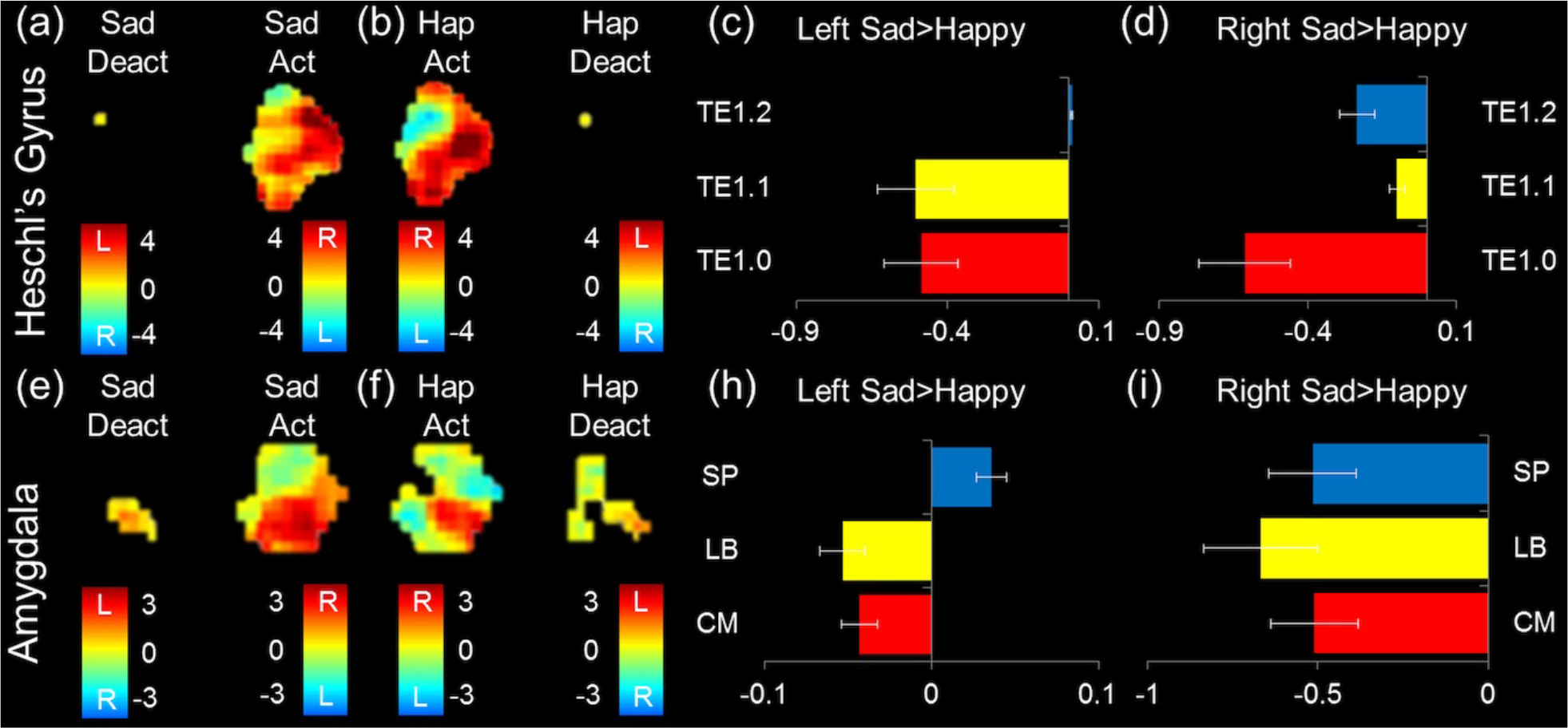
Heschl’s gyrus and amygdala activation and deactivation profiles with subdivision differences. Heschl’s gyrus (a) sad and (b) happy deactivation and activation with color bars coded for laterality (L/R). Sad > happy (c) left and (d) right hemisphere subdivision differences for activation. Amygdala (e) sad and (f) happy deactivation and activation with color bars coded for laterality (L/R). Sad > happy (c) left and (d) right hemisphere subdivision differences for activation.

## Discussion

We asked whether happy and sad emotions could be differentiated in HG and amygdala subdivisions [3]. We theorized a right hemispheric-leaning laterality would be associated with both happy and sad emotions [12–15]. The hypothesis was based on the right hemisphere favoring spectral cues [18]. Surprisingly, we found a right amygdala lateralization for sad whereas HG was left and right lateralized depending on subdivision. Both ROI responded more to sad than to happy. These findings suggest more research should be conducted to understand the hemispheric lateralization of emotion based on acoustic properties of the stimuli [18].

### Laterality subdivision differences for happy and sad emotion

A recent manuscript found a low-frequency pitch region towards the anterolateral HG Te1.2 [20] consistent with our results, where sad in minor tone showed greater activation within Te1.2 and both the left and right Te1.2 HG subdivisions responded more to sad than happy. Sad could evoke more activity from pitch responsive regions due to its tonality [1], its valence [3], or its dominance of spectral cues [18], however a previous study contrasting happy and sad music failed to find robust differences [8]. The study by Norman-Haignere et al. (2013) revealed a functional segmentation based on the same structural ROI utilized in the present manuscript, where a greater proportion of pitch sensitive voxels were found in the posterior-to-anterior gradient [20]. The current view of HG is that auditory fields fold across the rostral-caudal axis in a V-shaped gradient from high-to-low-to-high frequency [28]. The difference in encoding happy in a major key as rapid pitch changes and sad in a minor key as slow undulations could also be mirrored along this axis and lateralized by left and right hemisphere [18], not only in the spectro-temporal domain, but in valence of the stimuli (happiness and sadness or another contextualization of emotions: i.e. positive/negative).

The superficial amygdala was previously shown to be responsive to joyful music, while the laterobasal amygdala has been implicated in the evaluation of stimuli valence [3]. Differences in the functional lateralization of these subdivisions have not been previously evaluated in direct analyses [12–15]. Nevertheless, a meta-analysis has demonstrated that the amygdala was left lateralized in particular for negative emotions [14] and was primarily activated in the left hemisphere, centered in the superior and basolateral amygdala [14]. In the present study we found no differences within the left amygdala subdivision, possibility due to our stimuli set. However, we found significant differences between happy and sad stimuli within all subdivisions of the right amygdala activating more for sad, possibly due to the minor mode being more negative in valence [1, 3, 8, 10].

### Activation and deactivation subdivision differences

Previous studies of acoustic emotion have mostly concerned activation within HG and the amygdala [3, 11–15, 28]. In the present study we chose to separate activation and deactivation profiles within a subdivision [27]. This procedure along with the laterality calculations allowed us to determine the contributions of activation versus deactivation [27]. Activation profiles for HG mirrored previous laterality statistics with Te1.2 greater activation for sad compared with happy, with no significant deactivation notable. We would expect, due to the auditory nature of these stimuli, that HG would be considerably active bilaterally for both acoustic emotions, which is what was found. Previous studies have found right greater than left amygdala activation [1], but strong bilateral activation has been most consistently reported [3, 9]. Here we report slight differences in amygdala activation by subdivision contingent on which emotional stimuli (sad> happy). This finding is in line with the thought of sad emotions being negative in valence and activating the amygdala more strongly [3]. Amygdala deactivation has also been previously reported [15], but negative fMRI BOLD responses are currently under investigation for underlying causes [29]. Here we noted interesting patterns of amygdala deactivation (Fig. 1d, 2e and 2f); however, profiles failed to reach significance in statistical testing.

### Study Limitations and future directions

The present study asked if happy and sad emotion stimuli could be differentiated in HG and amygdala subdivisions based on an ROI hemispheric lateralization. Heschl’s gyrus is a known asymmetrical (left ≠ right) anatomical structure [26]; therefore we utilized hemispheric lateralization statistics [27] with cytoarchitectonic-MRI probabilistic maps of HG [26]. For area right (R) or left (L) asymmetrical and transformed to a symmetrical template, not included in the contralateral side, some issue may occur whereby its contribution is less or more than the nonincluded ipsilateral side depending on the activation or deactivation excluded/included due to the asymmetry. For example; areas right (R) and left (L), if area L not included in R, when computing the matrix absolute subtraction L from R within a symmetrical map (ss), (L-R =Lss), resultant Lss still contains L within the left side (where not in R) and now contributes more if activated, less if deactivated to the functionality of that side. Hence the contribution to the symmetrical side after transformation is more or less depending on areas included or excluded. This has the drawl back that the specific asymmetries may make lopsided results, if the asymmetry contributes predominately to the activation/deactivation profiles. The total original voxel space of left HG was 2,819 and right HG was 2,195 based on the 91-109-91 matrix, a difference of 22.14% in ROI area; therefore based on area would be biased left over right. Here we found significant results both left and right leaning, indicating due to the preponderance of voxel space on the left side, the right side for those results was considerably more active. The total original voxel space of left amygdala was 1, 831 and right amygdala 1,873, a difference of 2.24%, likely non-significant. Future studies should determine the exact inclusion or exclusion of the known anatomical asymmetries to hemispheric lateralization of emotion.

The present manuscript used 3 second stimuli from an emotion identification task that has previously categorized acoustic stimuli as happy or sad [16, 17]. Happy and sad acoustic stimuli have differing contributions of fine structure and envelope, which we theorized convey the emotion in sound. We hypothesized that happy and sad emotion could be differentiated by subdivision due to differing amounts of fine structure and envelope information contained within; however, it is quite possible that our pieces chosen from the emotion identification task [17], did not have adequate features to be distinguishable, hence why certain subdivision statistics failed to find significant differences. The observations in the present study due to hemispheric lateralization of emotion could be one of many attributes contributing to the identification of emotion such as distributed reward networks acting in synchrony [23]. Future studies should employ longer length excerpts in their original from mapped across time to determine differences in emotion [30].

## Conclusion

Our results demonstrate hemispheric laterality differences in HG and amygdala by subdivision. We found prominent right hemisphere responses in HG and amygdala subdivisions for sad but not happy. To some extent, changes in activation and deactivation profiles explained the extent of subdivision hemispheric laterality responses by emotion. Further studies should explore the laterality of activation and deactivation in emotional contexts.

## Acknowledgments

We thank the Posgrado en Ciencias Biomédicas, the Instituto de Neurobiología of the Universidad Nacional Autónoma de México (UNAM), the Consejo Nacional de Ciencia y Tecnología (CONACyT) México for the Graduate Fellowship 578458 to FAM Manno and CONACyT for the funding received via grant CB255462 to FA Barrios. We are grateful to L Casanova-Rico, A Orozco-Rivas for their administrative support; Z Gracia-Tabuenca, I Sánchez-Moncada, for their insightful comments; EH Pasaye and JJ Ortiz for their technical support; Dr. M.C. Jeziorski for his revision of the manuscript; Francis A. Manno Jr. and Helen D. Manno for all their support.

## Conflict of Interest

The authors declare no competing financial interests.

